# Evaluating WGS-based methods as an *in silico* replacement of phage typing for historical continuity and host association in *Salmonella* Typhimurium

**DOI:** 10.64898/2026.11.23.768569

**Authors:** George M. Kolodziejczyk, Jaromir Guzinski, Klara M. Wanelik, Mark Arnold, Alexessander Couto Alves, Liljana Petrovska, Jennifer M. Ritchie

**Affiliations:** School of Biosciences, Faculty of Health and Medical Sciences, University of Surrey, Guildford, Surrey, GU2 7XH, United Kingdom; Department of Bacteriology, Animal and Plant Health Agency (APHA – Weybridge), New Haw, Surrey, KT15 3NB, United Kingdom; Department of Epidemiological Sciences, Animal and Plant Health Agency (APHA – Weybridge), New Haw, Surrey, KT15 3NB, United Kingdom; School of Human Development, Faculty of Medicine, University of Southampton, Southampton, SO16 6HW, United Kingdom; Gastrointestinal Infections & Food Safety (One Health) Division, UK Health Security Agency, London, NW9 5EQ, United Kingdom

**Keywords:** *Salmonella* Typhimurium, subtyping, WGS, machine learning, host association, epidemiology, phage typing

## Abstract

2.

*Salmonella enterica* serovar Typhimurium (STm) poses a significant burden on public health worldwide. The Anderson phage typing scheme has historically supported surveillance, outbreak investigations and source attribution through associations between phage types and animal hosts. However, limitations inherent to the scheme have led to the gradual replacement of phage typing by whole-genome sequencing (WGS)-based methods. In this study, we applied statistical and machine learning approaches to 1,329 high-quality STm whole-genome sequences to compare phage typing with hierarchical clustering of core-genome multilocus sequence typing (HierCC) and the single nucleotide polymorphism (SNP) address. We observed low congruence (< 0.60) between phage typing and WGS-based approaches. Nevertheless, a random forest model using WGS-based subtypes as features was able to predict all phage types with high specificity (≥ 0.94) and at least 80% recall for 10 of the 24 analysed phage types. These included phage types of human and veterinary epidemiological importance, including DT193, DT104, DT2, DT40, DT56, DT9, and DT99. We then assessed whether WGS-based subtyping methods could replace the host-association feature of phage typing. At higher distance thresholds (≥ 100) WGS-based subtyping methods were more informative of animal host species and displayed stronger host-associations than phage types for 11 of the 12 animal hosts, with significantly stronger associations for ducks, finches, and pigs (all *p* < 0.05). Thus, despite low congruence between phage types and WGS-based subtypes, our random forest model accurately predicted several epidemiologically important phage types, including those important to public health and veterinary surveillance, supporting its potential as an *in silico* alternative for those phage types. Finally, our findings suggest HierCC and SNP address methods can capture the host-association information historically derived from phage typing.

**Impact statement:** Epidemiological tracing of isolates during outbreaks is central to effective public health outcomes. Whole-genome sequencing (WGS)-based approaches are increasingly replacing or supplementing traditional laboratory methods such as phage typing and serotyping, and low-resolution molecular approaches such as multi-locus sequence typing. Yet, for pathogens such as *Salmonella enterica* serovar Typhimurium (STm), these legacy methods underpin a substantial body of epidemiological knowledge that risks being lost in the transition to WGS-based surveillance. By applying two WGS-based subtyping methods to high-quality whole-genome sequences from 1,329 STm isolates, we demonstrate how WGS data can be leveraged to preserve and extend this knowledge. Despite limited congruence between phage types and WGS-based subtypes, a random forest model correctly predicted several epidemiologically relevant phage types, enabling continuity with historical phenotypic phage-typing data. In parallel, WGS-based methods show potential to capture animal host-associations as effectively as phage typing. Together, these findings show that WGS-based approaches can both bridge legacy phage-typing frameworks and inform upon host-association, reinforcing their value for national surveillance systems, government agencies and reference laboratories.

## 4. Introduction

Non-typhoidal *Salmonella* poses a significant public health burden globally, causing over 90 million infections and 84,000 deaths per year (1,2). In the United Kingdom (3,4), Europe (5), and the United States (6), most human salmonellosis cases are attributable to *Salmonella enterica* serovar Typhimurium (STm) and its monophasic variants. STm can infect a wide range of animal host species and is thus characterised as a generalist serovar of *Salmonella.* Bidirectional transmission between animals and humans facilitates both national and international spread of the pathogen (7). The substantial disease burden, associated mortality, and zoonotic potential of STm underscore the need for robust routine surveillance and effective source attribution.

Phage typing has been used for decades to subtype bacteria based on their susceptibility to lysis by defined panels of reference phages. For STm, the Anderson phage typing scheme, first described in 1976 (8), was considered the gold standard method of subtyping (9). The scheme utilises a panel of 34 typing phages and recognises 382 different phage types (8), comprising definitive types (DT) and “untypable” types (U), the latter of which are provisionally assigned (8). Despite nearly 50 years of use (8), the biological mechanisms underpinning phage type designations remain incompletely understood.

Phage typing has long supported routine surveillance, outbreak investigations, and characterisation of STm epidemiology (10–12). The utility of the phage typing scheme extends beyond subtyping to include host-association (10,13). It has been suggested that circulation of STm strains in specific animal hosts can differentially influence bacterial resistance and susceptibility to the Anderson typing phages (14). As such, phage typing may be uniquely informative of the primary animal host of some STm lineages (14). For example, DT2 and DT99 are associated with pigeons (15,16) whereas DT193 and DT104 exhibit a broader host range but are typically associated with pigs and cattle, respectively (12,17). Both host-adapted variants of STm (13,18) and successive waves of multi-drug resistant (MDR) STm clones that emerge approximately every 10 years (12), have been captured by phage typing. Monophasic STm DT193 is the dominant MDR clone currently circulating in the United Kingdom (12,19). Collectively, these features make the phage typing scheme valuable not only for routine surveillance but also for inferring host association and tracing the emergence and spread of epidemiologically important STm clones (20,21).

However, STm phage typing suffers from several limitations. Phage type switching, which occurs through the acquisition or loss of prophages and plasmids in STm, can undermine subtype stability and complicate epidemiological interpretations (14,22–24). The method also relies on subjective visual interpretation of lysis patterns, which has led to inconsistencies in phage type designations across laboratories (25–27). Additionally, reference phages can be difficult to propagate, primarily due to recombination with resident prophages (9,25), raising concerns that reference phage stocks held by national reference laboratories may become depleted in the near future. Consequently, several reference laboratories and government agencies have discontinued routine phage typing (20,22,28–30), creating the need and opportunity to implement whole-genome sequencing (WGS)-based subtyping methods for STm.

Widespread adoption of WGS has led to the replacement or supplementation of traditional phenotypic and molecular methods for characterising *Salmonella* (18,28,29,31–34). Two WGS-based subtyping methods widely used for routine surveillance and outbreak investigations are core-genome multilocus sequence typing (cgMLST) and single nucleotide polymorphism (SNP) address typing (28,35). cgMLST schemes utilise species specific cut-offs to define “core” loci, typically requiring that a locus be present in ≥ 95% of a set of representative isolates of the target microorganism. For example, the EnteroBase *Salmonella* cgMLST scheme comprises 3,002 loci present in ≥ 98% of 3,144 representative *Salmonella* isolates (18,36). Sequence variation at each locus defines distinct alleles, and the combination of alleles across loci yields an allelic profile for each isolate (18,37). Hierarchical clustering of cgMLST (HierCC) assigns isolates to clusters according to thresholds of allelic distance, defined as the number of differing alleles between isolates (38). This framework supports clustering at multiple levels of allelic difference (e.g. HC0, HC5, HC250), enabling epidemiological investigations from fine-scale outbreak detection to broadscale population-level analyses. The SNP address (39) identifies sequence polymorphisms by comparing the core-genome of each isolate against a reference genome and recording variant positions.

Single linkage clustering of isolates is performed at predefined pairwise SNP distance thresholds (0, 5, 10, 25, 50, 100, 250), generating a hierarchical ‘address’ that captures genetic relatedness at multiple levels of resolution (39). Genomic surveillance platforms such as EnteroBase, which supports cgMLST and HierCC-based subtyping, and SnapperDB, which implements the SNP address, have facilitated the subtyping of thousands of STm isolates for surveillance and epidemiological investigations (29,40,41).

The rapid growth of WGS technologies has made replacing laborious phenotypic methods such as phage typing, both feasible and timely. However, phage typing has historically supported outbreak investigations and source attribution of STm (25,30,42,43) and it remains unclear to what extent these features can be captured by WGS-based subtyping methods. The primary aim of this study was to determine whether WGS-based subtyping methods could replace or reliably predict phage type designations. The secondary aim was to assess whether these methods could replicate the host-associations captured by the phage typing scheme. To address these questions, we applied WGS-based subtyping methods to STm isolates collected during clinical investigations and routine veterinary surveillance in England and Wales. We assessed the discriminatory power and congruence of WGS-based subtypes relative to phage types and used a random forest model to evaluate the ability of WGS-based subtypes to predict phage type designations. Additionally, we compared the informativeness, completeness, and homogeneity of phage types and WGS-based subtypes with respect to animal host species and quantified the strength of their host-associations.

## 5. Methods

### 5.1 Strains, phage typing, and sequencing

The original study dataset comprised 1,586 STm isolates collected by the Animal and Plant Health Agency (APHA) through clinical investigations and routine surveillance in England and Wales between 2002 and 2023. Isolates were phage typed by trained APHA personnel. Isolates with: “no phage type” (NOPT), “untypable” (UNTY), “reacts but does not conform” (RDNC) or “not specified” phage type designations were excluded from the dataset. Metadata including the source of isolation (**Table S1**), year of isolation, sequence type (ST) based on the legacy seven-gene MLST scheme (44), serovar (**Table S2**), and phage type (**Table S3)** was collected for all isolates used in this study. Surveillance records were used to obtain the year and source of isolation; isolate serovars were confirmed phenotypically, via serotyping conforming to the White-Kauffman-Le Minor scheme (45), and *in silico* using an APHA in-house *Salmonella* typing pipeline.

The whole genome sequences of the STm isolates were generated as part of APHA surveillance activities, outbreak investigations, and research projects. DNA extraction, quality filtering of the short-read data, and *de-novo* genome assembly were performed as described in detail elsewhere (46).

### 5.2 WGS-based subtyping: SNP address generation

A SNP address was generated for each isolate from FASTQ files using SnapperDB v1.0.4 (39). The SNP address was separated into seven thresholds, corresponding to decreasing pairwise SNP distances: 250-SNP, 100-SNP, 50-SNP, 25-SNP, 10-SNP, 5-SNP and 0-SNP. For each threshold, a SNP address subtype represented a single-linkage cluster of isolates in which all pairwise SNP distances were less than or equal to the threshold. Higher-resolution thresholds (e.g., HC5) are expected to yield more subtypes each containing fewer isolates, than at lower-resolution thresholds (e.g., HC250).

### 5.3 WGS-based subtyping: scoring of cgMLST alleles

cgMLST alleles of the assembled STm genomes were called against the 3,002-locus *Salmonella* EnteroBase schema (May 2025 version) (18,33) using ChewBBACA v3.3.10 (37). The EnteroBase cgMLST schema was prepared using the “PrepExternalSchema” function, run with default parameters. Locus “STMMW_11461” was removed as no valid alleles were found for that locus within the study dataset. Invalid alleles were defined by chewBBACA as missing start or stop codons or containing stop codons in-frame. Subsequently, isolate cgMLST allelic profiles were derived using the “AlleleCall” function, run with default parameters. Alleles that were not an exact or highly similar match to loci in the schema were replaced with a value of 0 using the “ExtractCgMLST” function with the threshold parameter set to 0. Inferred alleles, defined as displaying less than 100% match to loci in the schema, were only imputed as the probable allele if they showed high similarity to loci within the schema (37). High allele similarity to loci in the schema was determined by chewBBACA using: a blast score ratio of 0.6 or greater, and allele length within 20% of the modal locus allele length (37). HierCC subtypes were derived from the cgMLST allelic profiles using HierCC v1.22 (34). Each subtype within a given threshold represented a cluster of isolates that exhibit an allele difference up to the value of the threshold (e.g isolates comprising an HC100 subtype, differ by no more than 100 alleles). To match the resolution and number of SNP address thresholds, only the HC0, HC5, HC10, HC25, HC50, HC100 and HC250 thresholds were examined for the HierCC subtyping method.

### 5.4 Discrimination index

The discriminatory ability of the subtyping methods was assessed by calculating the Hunter and Gaston discrimination index (DI) (47), a widely utilised index for the comparison of bacterial subtyping methods (31,48). The DI is defined as the ability to differentiate between distinct isolates (47), providing a measure of the resolution of a subtyping method. The DI and the associated 95% confidence interval (CI) (49) was calculated for each subtyping method (HierCC, SNP address, and phage typing). Significant differences were determined using the 95% CI of the DI of each subtyping method. The DI is shown in **Equation 1**.

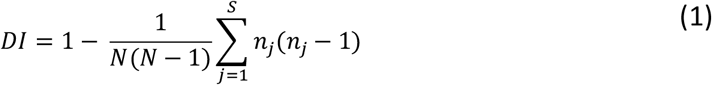

In **Equation 1**, *N* corresponds to the total number of subtypes within a subtyping method, and *n_j_* corresponds to the number of isolates belonging to the *j^th^* subtype.

### 5.5 Congruence between WGS-based subtyping methods and phage typing

Bidirectional congruence between phage typing and the WGS-based subtyping methods was assessed using the adjusted Wallace coefficient (AWC) (50). The AWC is defined as the probability of two isolates being classified into a single subtype by two different subtyping methods after correcting for chance agreement (50,51). The AWC has been used to assess the congruence between bacterial subtyping methods (48,52). An AWC value of 0 corresponds to chance agreement and values of 1 suggest perfect agreement between subtyping methods (50), representing a 100% probability of two isolates being assigned to a single subtype by both methods after correcting for chance agreement.

### 5.6 Input features, dataset split and machine learning algorithm selection

To reduce the risk of data leakage potentially arising from epidemiologically linked isolates, a non-clonal dataset was constructed (53–55). Data leakage can occur when highly similar isolates from a clonal group are present in both training and test sets leading to inflated accuracy estimates. Clonality was inferred when isolates shared five or fewer SNP differences (56) and were isolated from the same host and year (53–55). Isolates represented by phage types comprising fewer than five isolates were excluded to ensure sufficient examples for model training (**Table S3**). The non-clonal dataset, comprising one randomly sampled isolate per clonal group, was used for all model training and testing (**Fig. S1**). This dataset was randomly split into a training set and test set, comprising 80% (n = 598) and 20% (n = 150) of isolates in the dataset, respectively. The training and test sets were stratified by phage type, to avoid under-representation of the minority phage types.

Python v3.13.2 and the scikit-learn v1.6.1 (57) library was used for training and testing of all machine learning (ML) models in this study (**Table S4**). The three feature sets tested consisted of only the HierCC thresholds, only the SNP address thresholds, and both combined. Feature values comprised the subtypes of the different HierCC and SNP address thresholds. The “VarianceThreshold” function was used to assess whether invariant features were present in the training set and to remove such features, if found. No further feature selection was performed due to the low number of features in each dataset. Repeated stratified k-fold cross validation (RSKCV) was performed on the training set to identify the feature set and algorithm yielding the highest model predictive balanced accuracy across all algorithms tested (**Table S4**). The optimal algorithm for this dataset was determined by testing a selection of algorithms capable of multiclass classification. Each algorithm was run on the training set with default parameters and no fine-tuning using the three feature sets as inputs. A full list of algorithms tested can be found in **Table S4**. Balanced accuracy, a weighted accuracy that accounts for the potential differences in prevalence of each phage type in the dataset, was used to minimise overinflation of model performance due to imbalanced data. The random forest algorithm and the feature set comprising both HierCC and SNP address thresholds were chosen as they displayed the greatest balanced accuracy when compared to the other algorithms and feature sets tested (**Table S4**).

### 5.7 Random forest model hyperparameter tuning and testing

The random forest model was trained on the feature set comprising HierCC and SNP address thresholds. RSKCV, carried out on the training set with five splits and 10 repeats, was used to optimise the hyperparameters of the random forest model. Optimal hyperparameters were identified by evaluating balanced accuracy; the following model hyperparameters were identified as optimal: maximum features set to “none” and class weight set to “balanced_subsample”. Class (phage type) imbalance was addressed by weighting classes inversely proportionally to class size in the training dataset through the class weight hyperparameter (58). Subsequently, the prediction of phage types for isolates in the test set was performed using the random forest model with optimal hyperparameters.

### 5.8 Evaluation of random forest model performance on the test set

Tuned random forest model performance on the test set was determined using the precision, recall, specificity, F1-score, and a confusion matrix (**Fig. S2**) visualised using Matplotlib v3.10.7 (59). Additionally, overall balanced accuracy was used to evaluate model performance on the test set as it has been shown to be robust under class imbalance (60).

Spearman’s correlation coefficient (61,62), calculated using the “spearmanr” function in SciPy v1.15.2 (63), was used to determine whether a significant relationship existed between the number of isolates representing phage types in the training set and model F1-scores in the test set. Spearman’s correlation coefficient was chosen as the data deviated significantly from normality as assessed by a D’Agostino-Pearson test (64).

### 5.9 Feature importance analysis

The scikit-learn “permutation_importance” function (58) was used to identify the most important features for the prediction of phage types. The permutation importance function was run with 10 repeats on the test set.

Feature permutation involves randomly shuffling the values of a feature, effectively breaking the association between the feature and the label (phage type), and observing the decrease in accuracy score. The importance of each feature for the prediction of phage types was assessed using the median decrease in balanced accuracy score upon feature permutation. Balanced accuracy was used to ensure the feature importance was not biased towards the most represented phage types. Visualisation of the feature importance was performed using Matplotlib v.3.10.0.

### 5.10 Adjusted mutual information, homogeneity, and completeness of WGS-subtyping methods and phage typing with host groupings

For analyses involving host association, all isolates sampled from hosts represented by fewer than three isolates or from the environment were excluded (e.g “ENVIRONMENTAL_OTHER”) (**Fig. S1**).

To determine the similarity between phage typing and each WGS-subtyping threshold with animal host species, the adjusted mutual information (AMI), homogeneity, and completeness were calculated. The animal host species were treated as partitions where each distinct animal host was defined as a cluster of isolates sampled from that host. Animal host species were compared with phage types and WGS-based subtypes of each threshold. AMI assesses the agreement between a clustering (i.e., subtyping) method and a ground truth class (i.e., animal host species) and ranges from 0 (completely independent) to 1 (perfect agreement). AMI measures the reduction in uncertainty (entropy) in a variable when another variable is known, whilst correcting for chance agreement (65). Homogeneity measures whether each data point within a cluster shares the same class (animal host), and completeness evaluates whether all observations of the same class (animal host) are contained within the same cluster (WGS-based subtype or phage type) (66). AMI, homogeneity, and completeness were calculated using scikit-learn v1.8.0 (57).

The 95% CI of each metric was computed using jackknife resampling. Jackknife resampling was chosen over bootstrapping as this approach has been shown to yield 95% confidence intervals for clustering metrics that are robust to variation in the numbers of clusters (subtypes) and cluster size, and therefore more accurate (67).

### 5.11 Strength and significance of associations between phage types and WGS-based subtypes with specific animal hosts

Firth logistic regression (68), computed using firthmodels v0.7.2 (69) was used as to identify significantly host-associated phage types and WGS-based subtypes. Firth regression was used to reduce small-sample bias and account for perfect separation (70) between some phage types or WGS-based subtypes and animal hosts. Firth regression was run independently for each unique phage type and WGS-based subtype host combination across all WGS-based subtyping thresholds. Model *p*-values were computed using the likelihood ratio test based on the profile likelihood.

The phi coefficient was used to determine the strength of association for each of the host-associated phage types and WGS-based subtypes. 95% confidence intervals were calculated using bootstrap resampling. The Wilcox percentile bootstrap method for dependent correlations (71) was used to assess whether there was a significant difference in the strength of association between the most strongly host-associated phage types and WGS-based subtypes for each animal host species. The most strongly host-associated subtype was used for the comparison between methods as epidemiological investigations typically rely on a single strongly host-associated subtype per host (e.g DT99 in pigeons) (16,72).

Multiple comparisons correction was applied to the Firth regression and the Wilcox method bootstrap *p*-values using the Benjamini-Hochberg method (73).

## 6. Results

Of the original 1,586 isolates, 257 were excluded due to non-informative phage type designations (“no phage type” (NOPT), “untypable” (UNTY), “reacts but does not conform” (RDNC) or “not specified”). The remaining 1,329 STm isolates are hereafter referred to as the study dataset (**Fig. S1**).

### 6.1 Distribution of STm subtypes and source of isolation

The study dataset comprised STm isolates sampled from 24 different animal host species and six environmental or non-species-specific sources including “mammal”, “carnivore” and “birds” (**Table S1**). Among the STm isolates sampled from a species-specific host, 298 (26.4%) were from pigs, 236 (20.9%) from cattle, 224 (19.9%) from dogs, 88 (7.8%) from chickens, and 56 (5.0%) from horses. Twelve of the animal hosts were represented by no more than two isolates (**Table S1**). The remaining isolates originated from animal feed (177; 13.3% of the study dataset), environmental (16; 1.4%), and non-species-specific animal hosts (4; 0.3%). STm isolates within the dataset belonged to six legacy seven-gene MLST sequence types (ST19, 34, 568, 128, 313, and 323) and included 304 monophasic (22.9%) and 1,025 biphasic (77.1%) STm variants (**Table S2**).

**Table S5** summarises the number of subtypes and mean number of isolates per subtype generated by each subtyping method. Over increasingly stringent clustering levels (from HC250/250-SNP to HC0/0-SNP), HierCC and SNP address generated 19 (HC250) to 1,302 (HC0) and 27 (250-SNP) to 1,052 (0-SNP) subtypes, respectively. Accordingly, the mean number of isolates per subtype decreased. In comparison, phage typing resulted in 47 phage types, with the number of isolates per phage type ranging from 1 (DT5) to 368 (DT193) (**Table S3)**.

### 6.2 Discrimination index of the WGS-based subtyping methods and comparison to phage typing

Using phage typing as the reference comparator, we assessed the discriminatory power of each HierCC and SNP address threshold using the DI. Phage typing and all WGS-based thresholds showed high discriminatory power (DI > 0.8; **Table S6**), with the DI increasing at low HierCC and SNP address thresholds, consistent with their greater resolution (**Fig. 1**). Phage typing had a DI of 0.90 (95% CI [0.88, 0.91]). The DIs of all WGS-based thresholds differed significantly from that of phage typing except HC25 (0.91, 95% CI [0.90, 0.91]), HC50 (0.87, 95% CI [0.86, 0.88]) and 50-SNP (0.89, 95% CI [0.89, 0.90]). Of these, HC25 and the 50-SNP thresholds had DIs numerically closest to that of phage typing (**Fig. 1**).

**Fig. 1.**
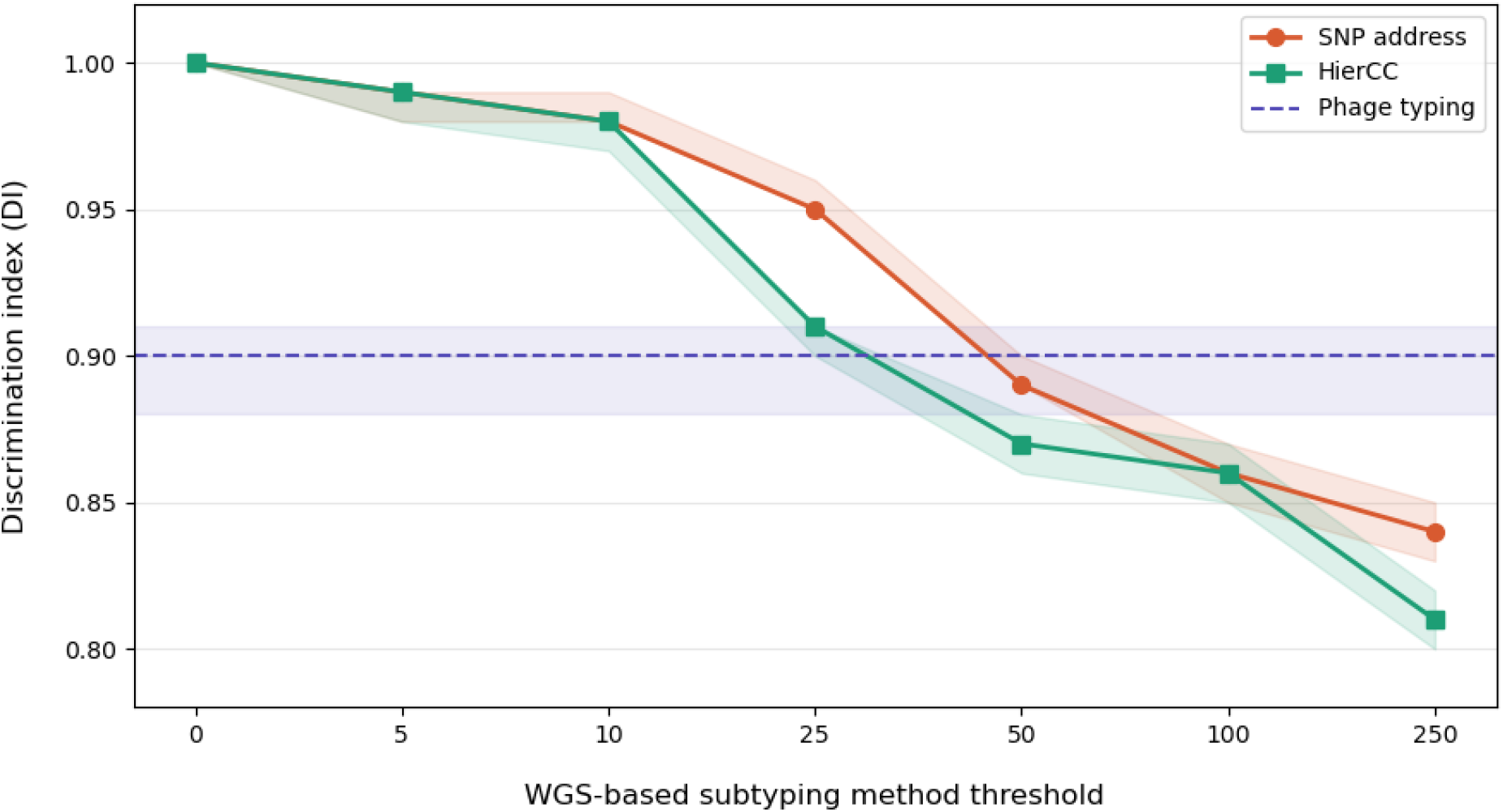
The Hunter and Gaston discrimination index (DI) of phage typing and two WGS-based subtyping methods: HierCC and SNP address, in a dataset of 1,329 *Salmonella* Typhimurium isolates. The x-axis represents increasing WGS-based subtyping thresholds (0–250). The y-axis (starting at 0.8) represents the DI value. The DIs of the phage typing, HierCC, and SNP address methods across thresholds are shown in purple, green, and orange, respectively. Solid shapes represent the DI values, with shaded regions representing 95% confidence intervals for each subtyping method. The DI value for phage typing is represented by the dotted line.

### 6.3 Congruence between WGS-based subtyping methods and phage typing

Congruence between WGS-based subtyping methods and phage typing was assessed using the AWC. Overall, congruence between WGS-based methods and phage typing (A vs B in Table 1) was low (AWC ≤ 0.6), indicating a low probability that two isolates classified in the same WGS-based subtype would share the same phage type. WGS-based thresholds 0-SNP and HC0 showed the highest congruence with phage typing (Table 1). In the reverse direction (phage typing compared to WGS-based subtyping; B vs A in Table 1) congruence decreased at higher-resolution thresholds and was greatest at thresholds ≥ 100, indicating that individual phage types were distributed among multiple WGS-based subtypes at these high-resolution thresholds.

**Table 1.** Bidirectional congruence (A → B and B → A, 95% CI) between each WGS-based subtyping method threshold (A) and phage typing (B) in *Salmonella* Typhimurium expressed as adjusted Wallace coefficients (AWC).

|  | **Phage typing (B)** |  |  |  |
|---|---|---|---|---|
| **WGS-based subtyping method threshold (A)** | **AWC<sub>(A vs B)</sub>** | **95 % confidence interval<sub>(A vs B)</sub>** | **AWC (<sub>B vs A)</sub>** | **95% confidence interval<sub>(B vs A)</sub>** |
| HC0 | 0.44 | 0.28-0.60 | 0.00 | 0.00-0.00 |
| HC5 | 0.35 | 0.31-0.39 | 0.04 | 0.03-0.05 |
| HC10 | 0.27 | 0.24-0.30 | 0.06 | 0.05-0.07 |
| HC25 | 0.40 | 0.37-0.43 | 0.36 | 0.31-0.41 |
| HC50 | 0.40 | 0.36-0.44 | 0.53 | 0.47-0.58 |
| HC100 | 0.39 | 0.35-0.43 | 0.54 | 0.49-0.60 |
| HC250 | 0.26 | 0.23-0.29 | 0.54 | 0.48-0.60 |
| 0-SNP | 0.60 | 0.54-0.65 | 0.01 | 0.00-0.01 |
| 5-SNP | 0.36 | 0.32-0.40 | 0.05 | 0.04-0.05 |
| 10-SNP | 0.36 | 0.33-0.40 | 0.06 | 0.05-0.06 |
| 25-SNP | 0.42 | 0.39-0.46 | 0.17 | 0.14-0.20 |
| 50-SNP | 0.41 | 0.38-0.43 | 0.41 | 0.36-0.46 |
| 100-SNP | 0.40 | 0.36-0.44 | 0.54 | 0.48-0.59 |
| 250-SNP | 0.34 | 0.31-0.37 | 0.55 | 0.49-0.61 |

### 6.4 Random forest model predictions of isolate phage types

Of the 1,329 isolates within our dataset, 785 comprised the non-clonal dataset (**Table S5**). After excluding phage types represented by fewer than five isolates (**Table S3**), 748 non-clonal isolates comprising 24 phage types remained and was used as the dataset for model selection, training, and evaluation on the test set (**Fig. S1**).

Multiple classification models were evaluated on the training set with RSKCV to determine the optimal algorithm and feature set to predict the phage types of the analysed isolates. Three feature sets were evaluated i) HierCC thresholds, ii) SNP address thresholds, and iii) a combined set of HierCC and SNP address thresholds, comprising seven, seven, and 14 features, respectively. None of the feature sets contained invariant features. Across all three feature sets, the random forest model achieved the highest balanced accuracy (**Table S4**) and was thus selected for subsequent modelling. The mean balanced accuracy for the predictions of the random forest model trained on the different feature sets and evaluated on the training set were: HierCC thresholds (46.6%, SD = 4.8%), SNP address thresholds (50.8%, SD = 5.6%), and the combined HierCC and SNP address thresholds (55.4%, SD = 4.8%). As the model trained on the latter showed the highest balanced accuracy, it was chosen as the feature set for model hyperparameter optimisation and evaluation on the test set.

Precision, recall, and F1 score of the model predictions of phage types for isolates in the test set are presented in Table 2. The model exhibited an overall balanced accuracy of 66.2%, when predicting the phage types of STm isolates in the test set, and achieved F1-scores ≥ 70% for nearly half (11/24, 45.8%) of the phage types. The random forest model exhibited ≥ 80% recall when predicting phage types DT105, DT116, DT193, DT2, DT41, DT56, DT9, and DT99 (**Fig. S2**). All phage types in the test set were predicted by the model with ≥ 94% specificity, indicating a low false positive rate for each phage type.

**Table 2.** Precision, recall, F1-score and sample size for each phage type predicted by a random forest model using 150 test set *Salmonella* Typhimurium isolates.

| **Phage type** | **Precision**<sup>\*</sup> | **Recall**<sup>†</sup> | **Specificity**<sup>‡</sup> | **F1-score**<sup>§</sup> | **Test set class sample size** |
|---|---|---|---|---|---|
| **DT1** | 0.33 | 0.40 | 0.97 | 0.36 | 5 |
| **DT104** | 0.67 | 0.77 | 0.96 | 0.71 | 13 |
| **DT104b** | 0.50 | 0.50 | 0.99 | 0.50 | 2 |
| **DT105** | 1.00 | 1.00 | 1.00 | 1.00 | 3 |
| **DT116** | 1.00 | 1.00 | 1.00 | 1.00 | 2 |
| **DT12** | 0.33 | 0.25 | 0.99 | 0.29 | 4 |
| **DT120** | 0.50 | 0.50 | 0.99 | 0.50 | 4 |
| **DT193** | 0.95 | 0.84 | 0.98 | 0.89 | 49 |
| **DT2** | 0.83 | 0.91 | 0.99 | 0.87 | 11 |
| **DT30** | 0.67 | 0.67 | 0.99 | 0.67 | 3 |
| **DT32** | 0.00 | 0.00 | 0.99 | 0.00 | 3 |
| **DT40** | 0.67 | 0.80 | 0.99 | 0.73 | 5 |
| **DT41** | 1.00 | 0.60 | 1.00 | 0.75 | 5 |
| **DT49** | 0.50 | 1.00 | 0.99 | 0.67 | 1 |
| **DT56** | 1.00 | 1.00 | 1.00 | 1.00 | 3 |
| **DT75** | 0.33 | 0.50 | 0.99 | 0.40 | 2 |
| **DT8** | 0.67 | 0.50 | 0.99 | 0.57 | 4 |
| **DT9** | 1.00 | 1.00 | 1.00 | 1.00 | 2 |
| **DT99** | 0.86 | 1.00 | 0.99 | 0.92 | 6 |
| **U288** | 0.43 | 0.67 | 0.94 | 0.52 | 9 |
| **U302** | 0.50 | 0.25 | 0.99 | 0.33 | 4 |
| **U308** | 0.00 | 0.00 | 0.99 | 0.00 | 4 |
| **U308a** | 0.50 | 0.75 | 0.98 | 0.60 | 4 |
| **U311** | 0.67 | 1.00 | 0.99 | 0.80 | 2 |

\* Represents isolates correctly predicted as phage type **X** divided by the isolates correctly predicted as phage type **X** and falsely predicted as other phage types instead of phage type **X**.
† Represents isolates correctly predicted as a phage type **X** divided by the isolates correctly predicted as phage type **X** and incorrectly predicted as phage type **X**. Equivalent to the true positive rate
‡ Represents the number of isolates correctly not predicted as phage type **X** divided by the number of isolates correctly not predicted as phage type **X** and the number of isolates incorrectly predicted as phage type **X**. Equivalent to the true negative rate
§ The F1-score represents the harmonic mean of the precision and recall values for each phage type.

Phage types represented by fewer than 10 isolates in the training set were considered low sample size phage types and accounted for 20.8% of all phage types in the training set (**Table S3**). When predicting these low sample size phage types, our model exhibited higher mean test set F1-score (0.79) than for phage types represented by ≥ 10 training set isolates (0.58). However, sample size for each phage type in the training set showed a weak positive, non-significant correlation with the test set F1-scores (r_s_ = 0.06, *p* = 0.77), indicating that training set sample size did not significantly impact the predictive performance of the model in this analysis.

### 6.5 WGS-based subtyping thresholds predictive of STm phage type

Feature importance was assessed using permutation importance, calculated using the median decrease in balanced accuracy for the test set predictions following repeated permutation of each feature. All WGS-based subtyping features (thresholds) contributed to the accurate prediction of STm phage types. Among them, 250-SNP showed the largest decrease in median balanced accuracy upon permutation (**Fig. 2**) and was therefore the most important feature for model predictions of phage type. The five most important features were, in decreasing order: 250-SNP, HC100, HC0, 50-SNP and HC25 (**Fig. 2**), indicating that both high (HC0) and low-resolution thresholds (e.g. 250-SNP) were important for accurately predicting phage types.

**Fig. 2.**
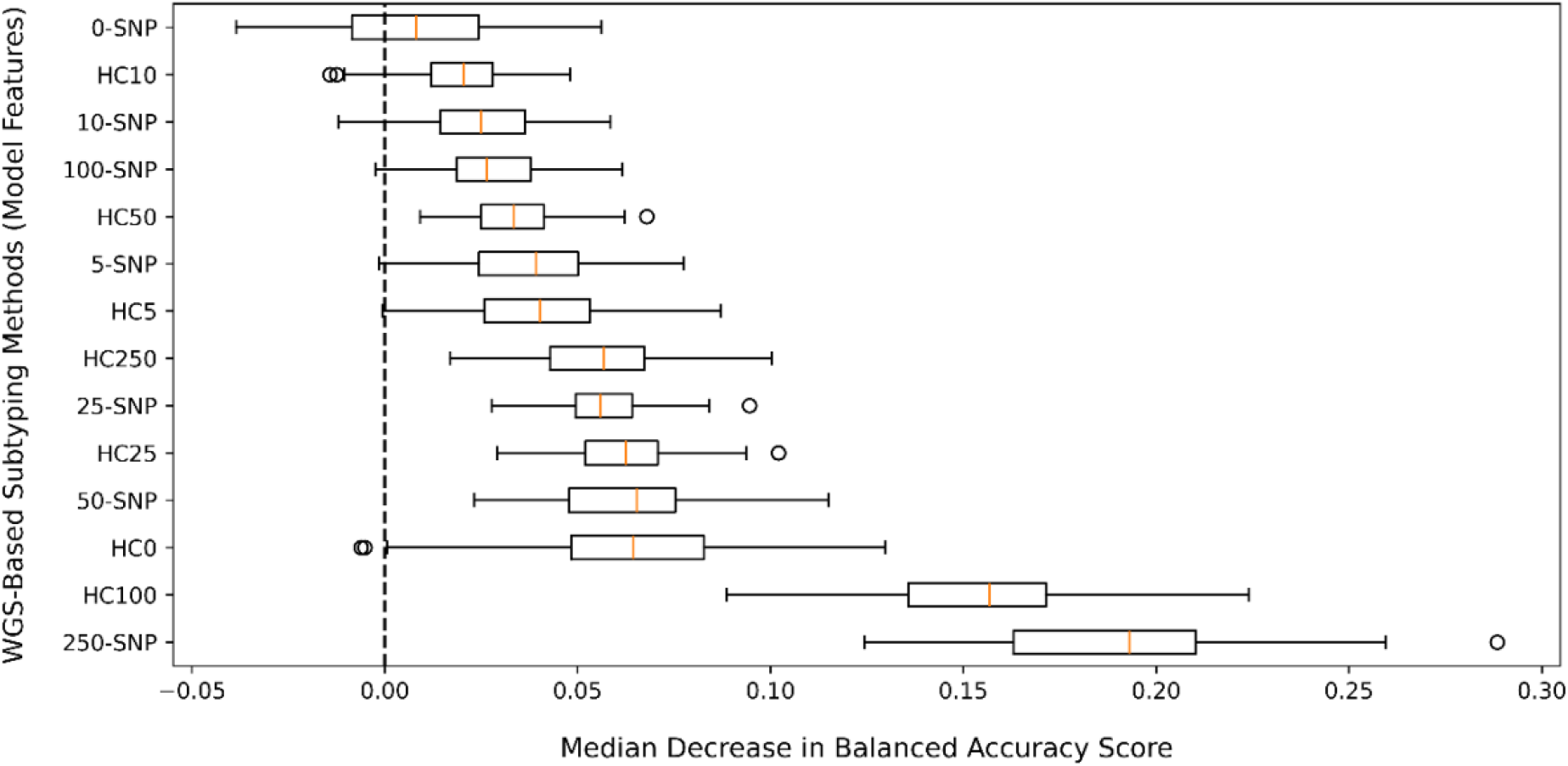
Permutation-based feature importance of WGS-based subtyping thresholds for predicting *Salmonella* Typhimurium phage types using a random forest model on 150 test set isolates. The figure shows the decrease in test-set balanced accuracy of the random forest model for phage type prediction following permutation of each WGS-based subtyping threshold (feature). The x-axis indicates the reduction in balanced accuracy score relative to the baseline balanced accuracy of the model (vertical dashed line). The y-axis lists WGS-based subtyping thresholds (features), ordered in ascending order of importance (top to bottom), according to the median decrease in balanced accuracy score. Boxes represent the interquartile range (IQR), with the orange line indicating the median. Whiskers extend to the most extreme datapoint within 1.5x the IQR, with points (dots) beyond this range plotted as outliers.

### 6.6 Agreement between phage types and WGS-based subtypes with animal host species

To evaluate agreement between subtyping methods and animal host species, we calculated the adjusted mutual information (AMI), homogeneity, and completeness for phage typing and each WGS-based subtyping threshold. After exclusion of isolates in the non-clonal dataset sampled from: a host represented by fewer than three isolates or isolates sampled from environmental sources, 661 isolates remained and formed the dataset used for host-association analyses (**Fig. S1**) (**Table S7**).

Across all WGS-based subtyping thresholds and phage typing, agreement with animal host species was low (**Table S8)**. The highest AMI scores among the WGS-based thresholds were observed for HC100 (0.421, 95% CI [0.382, 0.460]), 250-SNP (0.422, 95% CI [0.382, 0.461]), and 100-SNP (0.402, 95% CI [0.365, 0.439]) (**Fig. 3**). These thresholds had greater similarity to the animal host species than phage typing (0.352, 95% CI [0.319, 0.386]), indicating that their subtype designations were more informative of host species.

**Fig. 3.**
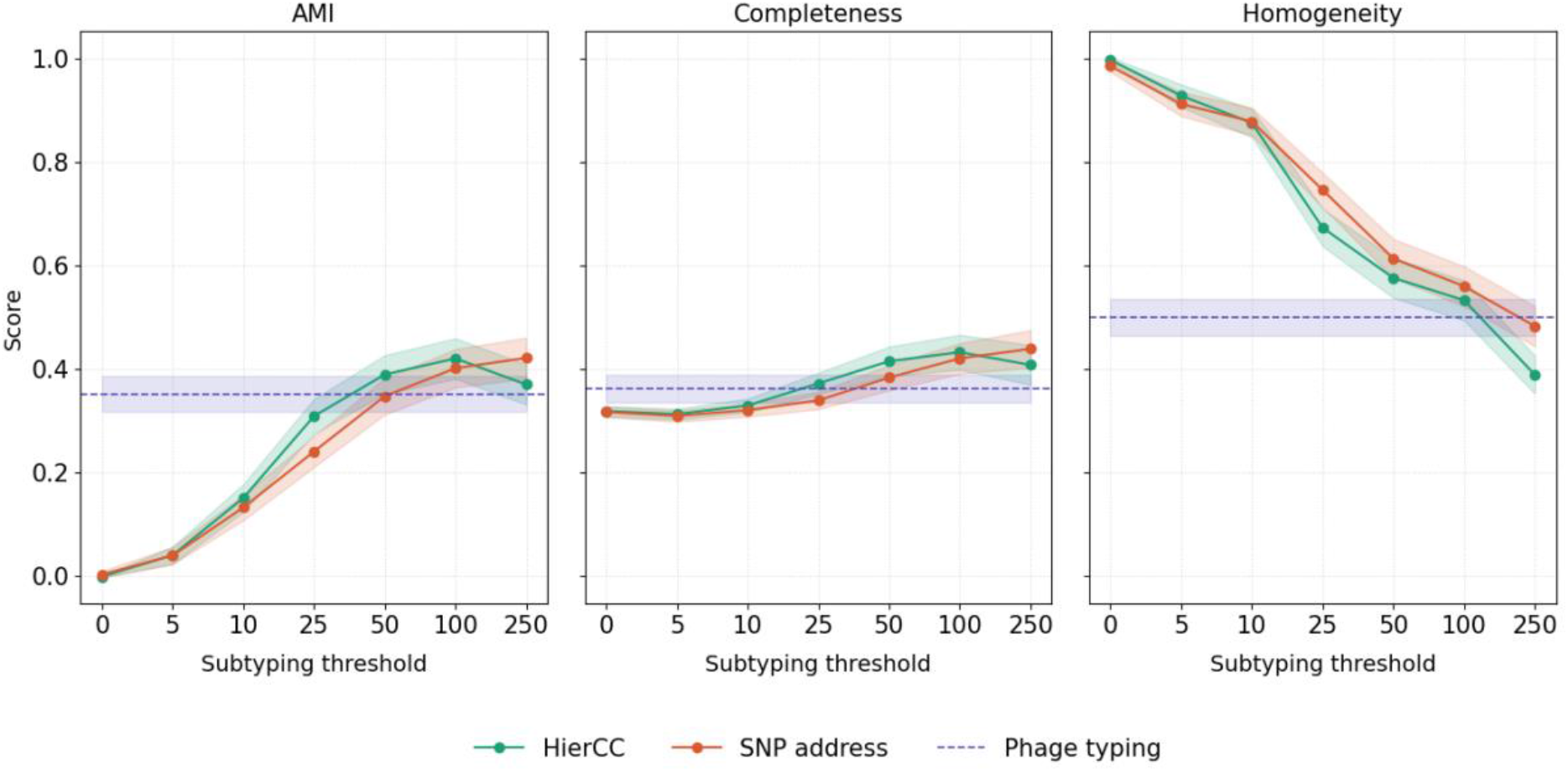
Adjusted mutual information (AMI), completeness, and homogeneity between animal host species and subtypes generated by HierCC and SNP address WGS-based subtyping methods and phage typing for *Salmonella* Typhimurium isolates. HierCC thresholds (green), SNP address thresholds (orange) and phage typing (purple) based on a dataset of 661 non-clonal *Salmonella* Typhimurium isolates. Points represent estimated scores and shaded regions represent 95% confidence intervals for each subtyping method.

At high resolution WGS-based thresholds (≤ 25), AMI was low (0.3) despite high homogeneity (> 0.80) (**Fig. 3**), indicating that these subtypes typically comprised isolates from the same animal host species. However, their low completeness score (< 0.35) suggested that isolates sampled from an animal host species were distributed across many different subtypes within these thresholds (**Fig. 3**). High homogeneity was driven primarily by many subtypes comprising few isolates at these thresholds. At low resolution WGS-based thresholds (≥ 50), all thresholds except 50-SNP, exhibited greater AMI scores than phage typing (**Fig. 3**). All WGS-based subtyping thresholds ≥ 50 also had higher completeness than phage typing, suggesting that at these thresholds isolates from the same animal host species were split across fewer WGS-based subtypes than phage types.

### 6.7 Comparison of the strength and significance of subtype-host associations between WGS-based subtypes and phage types

Significant associations between subtypes and host species were identified using the profile-likelihood derived *p*-values from Firth logistic regression models (**Tables S9-S11).** Across 12 animal hosts and different WGS-based thresholds, 59 SNP address subtype host associations, 33 HierCC subtype host associations, and 21 phage type host associations were statistically significant. The proportion of subtypes significantly associated with at least one host varied across the WGS-thresholds. The highest proportion occurred at lowest resolution WGS-based thresholds: 250-SNP (12/24, 50.0%) and HC250 (9/14, 64.3%), whereas much lower proportions occurred at highest resolution thresholds: HC0 (0/658, 0%) and the 0-SNP (1/641, 0.2%). In comparison, 21/43 phage types (52.5%) were significantly associated with an animal host. No phage types or WGS-based subtypes exhibited significant associations with chickens in our dataset, therefore only 11 animal hosts were examined in subsequent analyses.

The strength of host-association was quantified using the phi coefficient (φ), with values greater than 0.5 classified as strong associations. The greatest number of strongly host-associated subtypes was observed for HierCC and SNP address thresholds ≥ 50 (six for each WGS-based subtyping method). Conversely, no strongly host-associated subtypes were found for HierCC and SNP address thresholds ≤ 5. Four phage types exhibited strong host-associations. For each host species, we compared the strongest association identified by each WGS-based method with the strongest phage type association. WGS-based subtypes showed stronger associations than phage types for 11 host species (**Fig. 4**), although the differences were only statistically significant for ducks (HierCC only; adjusted *p* < 0.01), finches (adjusted *p* < 0.05), and pigs (adjusted *p* < 0.0001). Phi coefficients for all WGS-based subtypes and phage types across all 11 animal hosts are reported in **Table S12**.

**Fig. 4.**
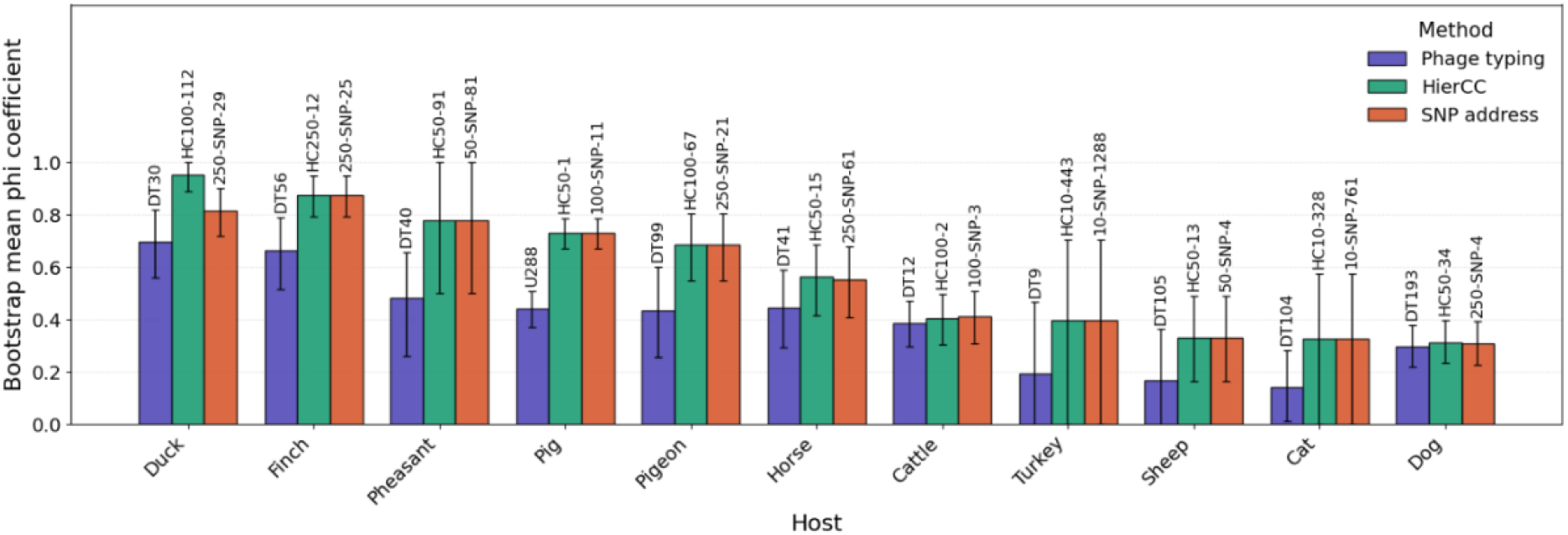
The strength of subtype-host associations for 661 non-clonal *Salmonella* Typhimurium isolates sampled from 11 animal host species based on WGS-based subtypes and phage types. Three subtyping methods were compared: phage typing (purple), HierCC (green), and SNP address (orange). For each of the 11 animal hosts, the WGS-based subtypes and phage types with the strongest host associations (adjusted *p* < 0.05), assessed using the phi coefficient, are shown above the bars. Bars represent the phi coefficient; error bars represent 95% confidence intervals.

## 7. Discussion

The transition to WGS-based subtyping risks loss of continuity with historic phage typing data because of the unknown relationship between these two subtyping approaches, and many historical isolates being unavailable for sequencing. Here, HC25 and 50-SNP thresholds provided similar discriminatory power as phage typing. Although directional congruence between WGS-based subtypes and phage types was low (AWC ≤ 0.6), a random forest model predicted several epidemiologically important phage types including DT193 and DT104, with F1-scores of at least 0.7. All WGS-based thresholds were important for model predictions of phage types (**Fig. 2**), permutation of the 250-SNP feature producing the greatest decrease in balanced accuracy. Lower-resolution WGS-based thresholds (≥ 50), except 50-SNP, were also more informative of animal host species than phage typing. At these thresholds isolates from the same animal host were grouped within fewer subtypes resulting in higher completeness. Conversely, higher-resolution thresholds (< 50) generated more host-homogeneous subtypes but distributed isolates from each host across more subtypes, resulting in lower completeness. For each of the 11 animal hosts examined, the strongest WGS-based subtype association exceeded the strongest phage type association, although this difference was only significant for ducks, finches, and pigs. These findings suggest that WGS-based subtyping can preserve and, for some hosts, enhance the capture of host-association signals previously inferred from phage types.

Although directional congruence between WGS-based subtypes and phage types was low (AWC ≤ 0.6), several factors may explain the low congruence between WGS-based methods and phage typing. Most notably, these two approaches capture different underlying biological processes. Phage typing reflects a whole-organism phenotype in response to a panel of lytic phages and can be influenced by the presence or loss of genetic elements (e.g. prophages (14), plasmids (74), CRISPR-associated systems (9)), as well as variation in surface structures and phage receptors during animal host carriage (75). The mechanism(s) that govern phage type designation in *Salmonella*, including the precise genetic determinants of phage susceptibility and resistance, remain incompletely understood (11,12). In contrast, HierCC and SNP address primarily index core-genome relatedness, with clustering thresholds optimised for genetic distance evaluation at varying scales rather than corresponding to phage-susceptibility patterns. Consistent with this, previous work has shown that WGS-based approaches frequently redefine *Salmonella* lineages created by phenotypic schemes (18,22,76).

Our random forest model, trained on a combination of HierCC and SNP address subtypes, correctly predicted nine of 24 phage types with high F1-score (≥ 0.70), suggesting a low rate of false-positive and false-negative predictions for those phage types. These included several phage types with human and veterinary epidemiological importance such as DT104 (77), DT193 (19,78), DT2 (16), DT40 (10,79,80), DT41 (81), DT9 (21,82), DT99 (15,16). Phage type DT104, and more recently DT193, represent dominant MDR STm clones that have been associated with numerous human outbreaks (26,77,78,83). In the veterinary context, DT2 and DT99 (15,16), as well as DT40 and DT56 (84,85), were associated with avian outbreaks, in pigeons and passerines, respectively. The ability to predict these phage types from WGS-based subtypes with a relatively high degree of accuracy using a machine learning modelling framework provides a practical means of maintaining continuity with historical phage typing data and supports long-term analyses of STm epidemiology as surveillance transitions to WGS-based frameworks.

A possible explanation for poor predictive performance of the random forest model for certain phage types is phage type switching. Phage types are known to undergo switching, where strains that are very closely related genetically can have different phage type designations, and can be polyphyletic, arising independently from different genomic backgrounds (22). In our test data set, 25% of DT120 isolates and 33% of DT30 isolates were incorrectly predicted by the model as U311 and DT8, respectively, consistent with previous reports of phage type switching between these pairs (14,24). Such switching is typically associated with the acquisition or loss of accessory genetic elements (14,74) and may occur without substantial changes in the core genome. Consequently, as our model relied on core-genome-based subtypes, it may have therefore failed to distinguish between isolates that differed in phage type primarily due to changes in the accessory genome. ML approaches that incorporate accessory-genome or whole-genome features such as prophage content, plasmid content, or virulence determinants, could improve discrimination between phage types known to undergo switching such as DT120/U311 and DT30/DT8 (14,24).

We noted several instances where feature importance for phage type prediction did not align with congruence between WGS-based subtypes and phage types. For example, 250-SNP was identified as the most informative feature for the random forest model predictions (**Fig. 2**) yet showed relatively low congruence with phage typing (AWC = 0.33; Table 1). Conversely, the 0-SNP threshold exhibited the highest congruence with phage types (AWC = 0.60) but was the least important feature for model predictive accuracy. This discrepancy between congruence and predictive importance likely reflects the complex relationship between phage types and genomic lineages. Many phage types are polyphyletic or have arisen independently in multiple genetic backgrounds (12,13,22), so lower-resolution thresholds may capture multiple phage types in a single WGS-based subtype, reducing overall congruence but providing broader-scale structure that is informative for random forest model predictions. In contrast, higher-resolution thresholds may group genetically related isolates that carry different phage type designations, for example due to phage typing switching (74), thereby diluting the predictive value of individual subtypes. Together, these observations emphasise that congruence of the evaluated WGS-based thresholds and subtypes with phage typing does not necessarily translate into useability of these subtypes for in silico phage type prediction.

Despite limited congruence with phage types, WGS-based subtyping methods captured host-association patterns at least as effectively as phage typing. Several WGS-based thresholds showed greater agreement with animal host species than phage typing, although the balance between completeness and homogeneity varied with resolution; lower-resolution thresholds had greater completeness, whereas higher-resolution thresholds produced more homogeneous but less complete subtypes. The strongest host-associated WGS-based subtypes were significantly stronger than the corresponding phage type designations for associations with ducks, finches and pigs, outperforming the well-established associations of DT30, DT56, and U288 with these hosts, respectively (14,87–89). Together, these patterns may suggest that differences in the strength of host-association between WGS-based subtypes and phage types likely reflect host-adapted STm lineages, better captured by core-genome (WGS)-based methods, than mechanisms underpinning phage type designation, such as prophage or plasmid acquisition (14,74).

The mechanism(s) that underlie phage type designation and their impact on host-association remain incompletely understood (12,14). It has been proposed that exposure to distinct phage communities within animal hosts imposes host-specific selective pressures on circulating STm lineages, favouring resistance profiles that contribute to the emergence of host-associated phage types (14). If correct, this hypothesis would imply that phage typing has an intrinsic advantage over purely genomic methods for host-association, as it directly captures variation in phage susceptibility. However, our data do not support this as a general pattern, as WGS-based subtypes at thresholds with discriminatory power comparable to or weaker than phage typing exhibited greater informativeness of animal host species. Furthermore, the most strongly host-associated WGS-based subtypes did not differ significantly in the strength of host-association when compared to phage types, with significant differences in favour of WGS-based subtypes in three of the 11 animal hosts examined. Instead, our findings support core-genome relatedness as an informative variable for host-association, consistent with previous work on host prediction for *Salmonella* isolates (53,86,87). The relative advantage of genetic relatedness over phage-based designations appears to be host-dependent, but in this study WGS-based subtyping outperformed or matched phage typing for all examined animal host species.

This study has several limitations. First, the dataset comprised 1,329 STm isolates representing 27 phage types and 24 animal host species, collected in England and Wales during clinical investigations and routine veterinary surveillance, which may limit the generalisability of our findings to other geographic regions, host populations, serovars, bacterial species, and phage-type designations. Sampling bias and differences in phage type and host prevalence (20) likely contributed to class imbalance, a common issue in ML classification problems (88,89). Second, although reasonable efforts were made to account for the potential of epidemiological linkage between isolates, such linkage might have persisted and modestly inflated accuracy estimates. Third, the outputs of the WGS-based subtyping methods depend on the underlying schema and reference genomes (39,90), and results may differ if alternative frameworks were used. Finally, the performance of our random forest model was constrained by the training data set (comprising 24 phage types), as phage types represented by fewer than five isolates were excluded; predictive accuracy for rare phage types therefore remains uncertain and will require evaluation in larger, more diverse datasets.

In conclusion, core-genome WGS-based subtyping methods are unlikely to provide a one-to-one replacement for phage typing as a classification scheme. Nevertheless, our findings show that they can preserve and at times enhance the epidemiological information historically derived from phage types. Across a range of thresholds, WGS-based subtypes captured host-association signals at least as well as phage typing and, at lower-resolution thresholds, proved more informative for host-association. Furthermore, our results demonstrated that WGS-based subtypes could be used within a random forest model to directly predict several epidemiologically relevant phage types with high recall and specificity, providing a practical bridge to legacy phage-typing data. Overall, and within the constraints of the study dataset, these results support the use of WGS-based methods as a viable replacement for phage typing for host-association analyses and source attribution, particularly in veterinary surveillance.

## 8. Author statements

### 8.1 Author contributions

G.M.K: conceptualisation, data curation, formal analysis, investigation, methodology, software, visualisation, writing – original draft, writing – review & editing.

J. G: conceptualisation, data curation, resources, supervision, writing – review & editing.

K.M.W: conceptualisation, supervision, writing – review & editing.

M. A: conceptualisation, supervision, writing – review & editing.

A.C.A: conceptualisation, funding acquisition, supervision, writing – review & editing.

L.P: conceptualisation, funding acquisition, supervision, writing – review & editing.

J.M.R: conceptualisation, funding acquisition, supervision, writing – review & editing.

### 8.2 Conflicts of interest

The authors declare that there are no conflicts of interest.

### 8.3 Funding information

G.M.K received PhD studentship funding from the University of Surrey Doctoral College and the Animal and Plant Health Agency.

### 8.4 Ethical approval

This study was assessed for ethical and governance issues using the University of Surrey’s ‘Self-assessment for governance and ethics – animal research (SAGE-AR)’ platform. The described activities did not meet the minimum threshold required for further review.

### 8.5 Consent for publication

### 8.6 Acknowledgements

## Notes

### Competing Interest Statement

The authors have declared no competing interest.

